# Cantilever Sensors for Rapid Optical Antimicrobial Sensitivity Testing

**DOI:** 10.1101/679399

**Authors:** Isabel Bennett, Alice Pyne, Rachel McKendry

## Abstract

Growing antimicrobial resistance (AMR) is a serious global threat to human health. Current methods to detect resistance include phenotypic antibiotic sensitivity testing (AST) which measures bacterial growth and is therefore hampered by slow time to result (~12-24 hours). Therefore new rapid phenotypic methods for AST are urgently needed. Nanomechanical cantilever sensors have recently shown promise for rapid AST but challenges of bacterial immobilization can lead to variable results. Herein a novel cantilever-based method is described for detecting phenotypic antibiotic resistance within ~45 minutes, capable of detecting single bacteria. This method does not require complex, variable bacterial immobilization, and instead uses the laser and detector system to detect single bacterial cells in media as they pass through the laser focus. This provides a simple read out of bacterial antibiotic resistance by detecting growth (resistant) or death (sensitive), much faster than current methods. The potential of this technique demonstrated by determining resistance in both lab and clinical strains of E. coli, a key species for clinically burdensome urinary tract infections. This work provides the basis for a simple and fast diagnostic tool to detect antibiotic resistance in bacteria, reducing the health and economic burdens of AMR.

## Main

Antimicrobial resistance (AMR) is steadily increasing and poses a major threat to global health, with estimates of AMR leading to 10 million deaths per year and costing the global economy $100tn by 2050.^[1, 2]^ The increase in AMR has been caused by several factors including the overuse of antibiotics.^[3]^ Despite the growth of AMR, methods for antibiotic susceptibility testing (AST) have remained relatively unchanged for several decades.

In common AST methods, bacterial growth is used as a measure of sensitivity to antibiotics, determined directly by an increase in media turbidity (the number of bacteria) or indirectly by the release of fluorescent metabolites. These phenotypic methods provide *in vitro* confirmation of resistances in isolated bacterial species, which are inferred from known resistance genes in genetic methods. However phenotypic methods are inherently limited by the speed of bacterial growth (for example, the doubling time of *E. coli* is 20 minutes, whereas *M. tuberculosis* is 15-12 hours), meaning these methods require long culture times (12-24 hours, or longer for some species) for an observable change to occur. These delays result in empirical prescribing of antibiotics for patients instead of targeted treatment, which has been shown to increase mortality from sepsis fivefold,^[4]^ in addition to being a driver of resistance. Having access to the identity and antibiogram of the pathogen just a few hours earlier could avoid unnecessary costs associated with inappropriate prescribing, increase patient welfare, and reduce the effects of AMR. ^[5, 6]^ Therefore to reduce the damaging effects of AMR, we require solutions in the form of novel diagnostic tools to detect resistance and improve antibiotic stewardship, surveillance and patient management. ^[7, 8]^

Recent developments in this field have exploited single cell methods for faster and more sensitive detection of antibiotic resistance. This has been achieved by miniaturizing the volume observed using microfluidics,^[9–11]^ measuring mass or mechanical changes,^[11–13]^ or by exploiting machine learning techniques for video tracking analysis of single cells.^[14–16]^ Despite advances in the detection limit, and speed of testing, these are mostly complex set-ups, which remain far from point of care.

Recently a nanomechanical method of detecting the viability of bacterial cells immobilized on soft cantilevers was reported by Longo *et al*^[17–20]^ which has attracted much attention by virtue of its ability to detect AST within minutes. In the nanomechanical method a non-specific linker molecule was used to coat the cantilever surface with hundreds of bacterial cells, and the motion of the cantilever was monitored using a laser before and after the application of an antibiotic. However there remains speculation as to the origin of the nanomechanical signal.

Herein when we initially sought to reproduce the Longo method and apply it to clinical samples, a significant issue was found in obtaining consistent bacterial immobilization on the cantilever surface. Bacterial immobilization numbers were found to vary from cantilever to cantilever, from zero / low numbers to very high clumpy immobilization (Figure S1). Efforts to identify the source of variability by testing different immobilization conditions resulted in significant variation across conditions (from 90 to >1000 cells, Figure S2) with no clear pattern identified. For example, of 60 cantilevers functionalized with bacterial cells, only 28 achieved measurable bacterial immobilization, of which only five had “optimal coverage” of 500-600 cells. Additionally, no significant difference was found between the mechanical cantilever motion pre-antibiotic and post-antibiotic treatment for these five cantilevers (P=0.4569, Figure S3). However, many large ‘peaks’ observed in the raw data were observed to correlate temporally and spatially with bacterial cells in solution passing through the laser path. Here we exploit this optical signal as an alternative method to the nanomechanical method reported by Longo *et al*, with the advantage that it does not require complex immobilization chemistries and optimization of bacterial seeding densities. This novel optical method is able to rapidly detect antibiotic resistance in bacterial solutions with single cell resolution.

Our method uses a cantilever, laser and sensitive photodetector to measure the effect of antibiotics on bacterial growth, as briefly described here. A reflective surface (small stiff cantilever) is immersed in filtered growth media, off which a laser is reflected onto a photodiode detector (**Figure 1**a). Stiff cantilevers (AC160 TS, *k* = 26 N/m) were selected to disentangle the effect of the optical signal from the nanomechanical motion of the cantilever. In bacterial growth media (LB media) before inoculation with bacterial cells, no variation in the laser signal was observed (Figure 1b). On inoculation with bacterial cells, the bacteria free in the growth media are observed to move through the path of the laser. This cell movement in solution interferes with the laser beam, causing it to shift on the detector, observable as peaks in the signal (Figure 1c). On addition of antibiotic to the media, cell death occurs in antibiotic-sensitive bacteria, and fewer bacteria are detected passing through the laser. This results in a decrease in the number of peaks after ~45 minutes (Figure 1d).

**Figure 1.**
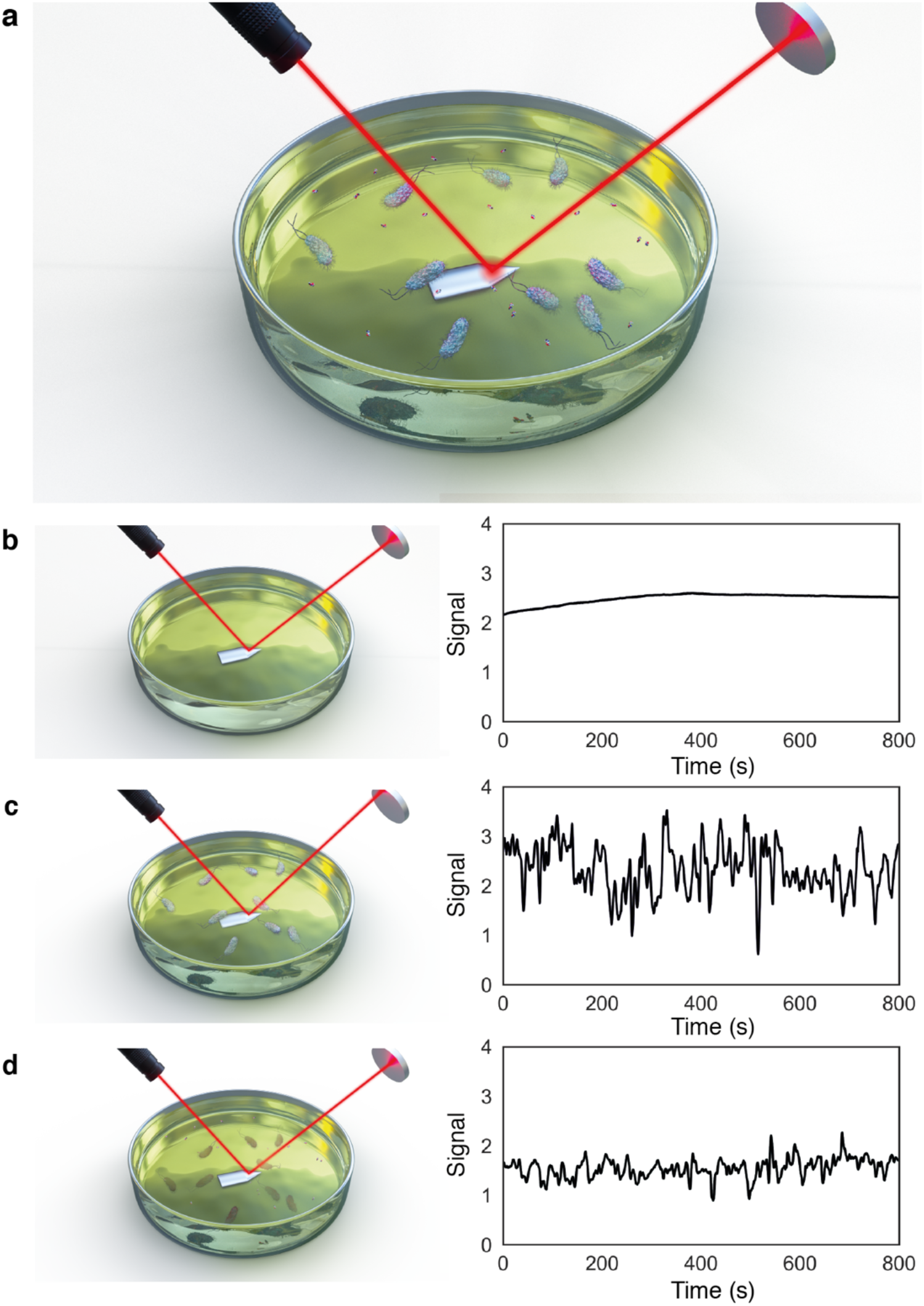
Principles of the rapid optical AST method. **a**, Illustration of bacterial cells inoculated in growth media with antibiotic molecules, with laser reflecting off cantilever surface onto photodiode detector. Bacteria in solution move through the laser beam, which can be observed as peaks in the photodiode signal. The photodiode signal measured in media solution decreases after addition of antibiotic for sensitive strains. **b**, **c**, **d**, Photodiode signal without bacterial inoculant (**b**), with bacteria in solution (**c**) and 45 mins after addition of antibiotic (**d**).

To determine the origin of the signal, the bacterial concentration in solution was reduced to ~ 10^5^ CFU (colony forming units, a standard measure of bacterial concentration). At this concentration individual bacterial crossing events can be observed as peaks within the optical signal (**Figure 2**a). When a single bacterium is tracked optically crossing the path of the laser (Figure 2b, blue circle), a corresponding peak in the signal can be observed in the data (Figure 2c). These peaks are of varying width and amplitude, due to differing angle and distance at which the bacteria pass through the laser. As more bacteria are added to the system (i.e. increasing CFU), the number of peaks in the signal also increases (Figure 2d), indicating that it is the bacteria giving rise to the signal.

**Figure 2.**
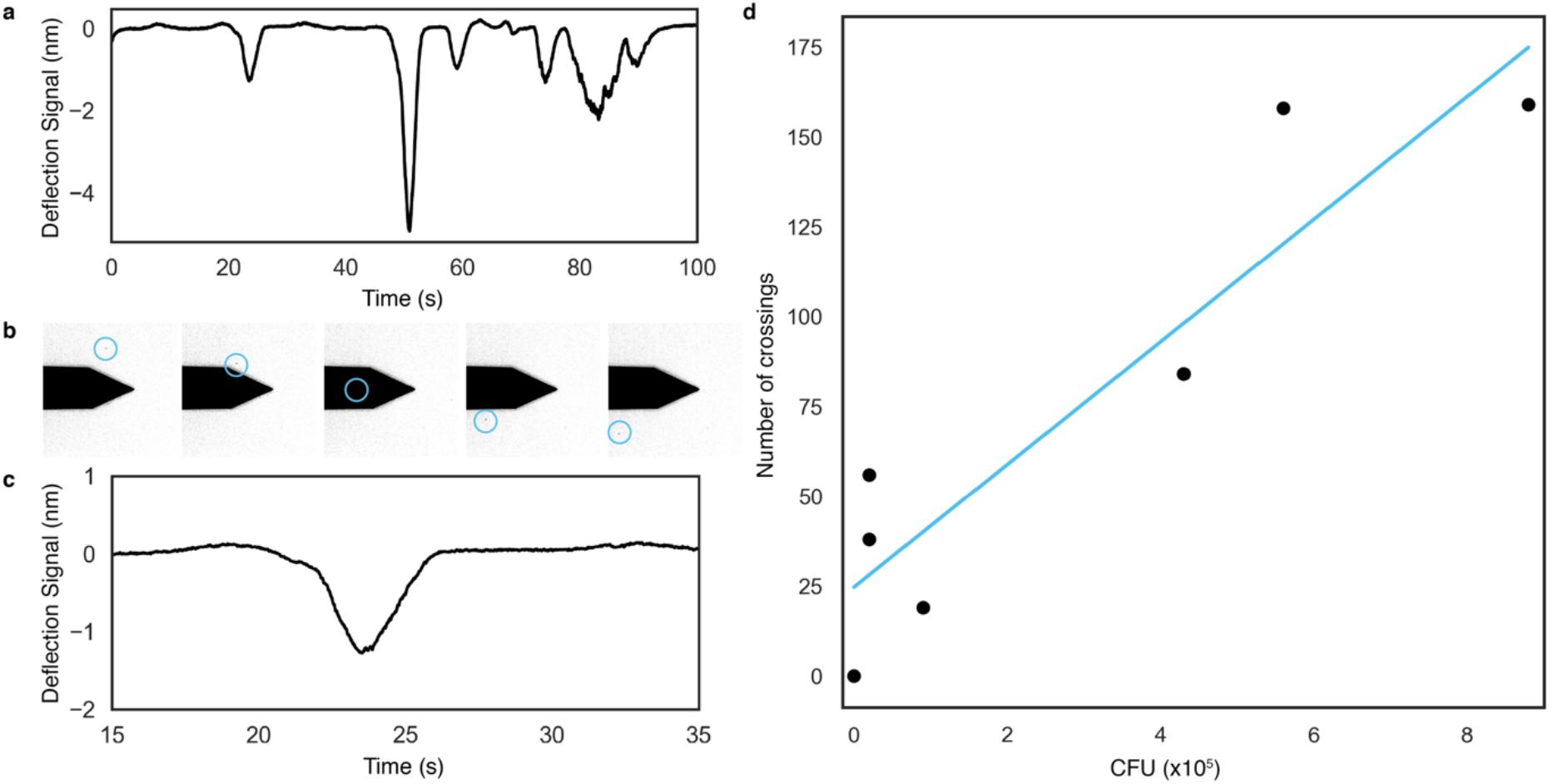
Signal caused by bacteria crossing the laser path decreases after 45 minutes from antibiotic addition. **a**, At low bacterial inoculant concentration, individual peaks can be identified within the signal. Combined optical tracking and signal measurement shows a of single bacterium (blue circle) passing through the laser path (**b**, optical images) as a single peak in the signal (**c**). **d**, Bacterial concentration (CFU, x10^5^) correlates with the number of bacterial crossings.

The number of peaks observed in the raw signal is linked to the number of viable bacteria in solution, which can exploited to determine antibiotic resistance. If the number of peaks (or bacterial crossings) is measured at distinct time points during an experiment (for example ‘media only’ (blue box), ‘inoculated media’ (green box), ‘inoculated media containing antibiotic’ (red box)) (Figure S4), a distinct trend appears where bacterial crossings increase on addition of bacteria to the system (Figure 3a, at blue dotted line), and decrease around 45 minutes (about two replication cycles for *E. coli*) after the addition of antibiotic (yellow dotted line) in the case of sensitive strains. The two peaks observed in the signal correspond to the addition of bacteria and antibiotic (**Figure 3**a, blue and yellow dotted lines, respectively) and occur due to mixing of the system. These peaks settle to a baseline and are observed in control experiments (Figure S5, points ‘3’ and ‘4’). This trend is not observed in a control where growth media is added without antibiotic, as the number of crossings continues to rise over time due to cell replication (Figure S5).

**Figure 3.**
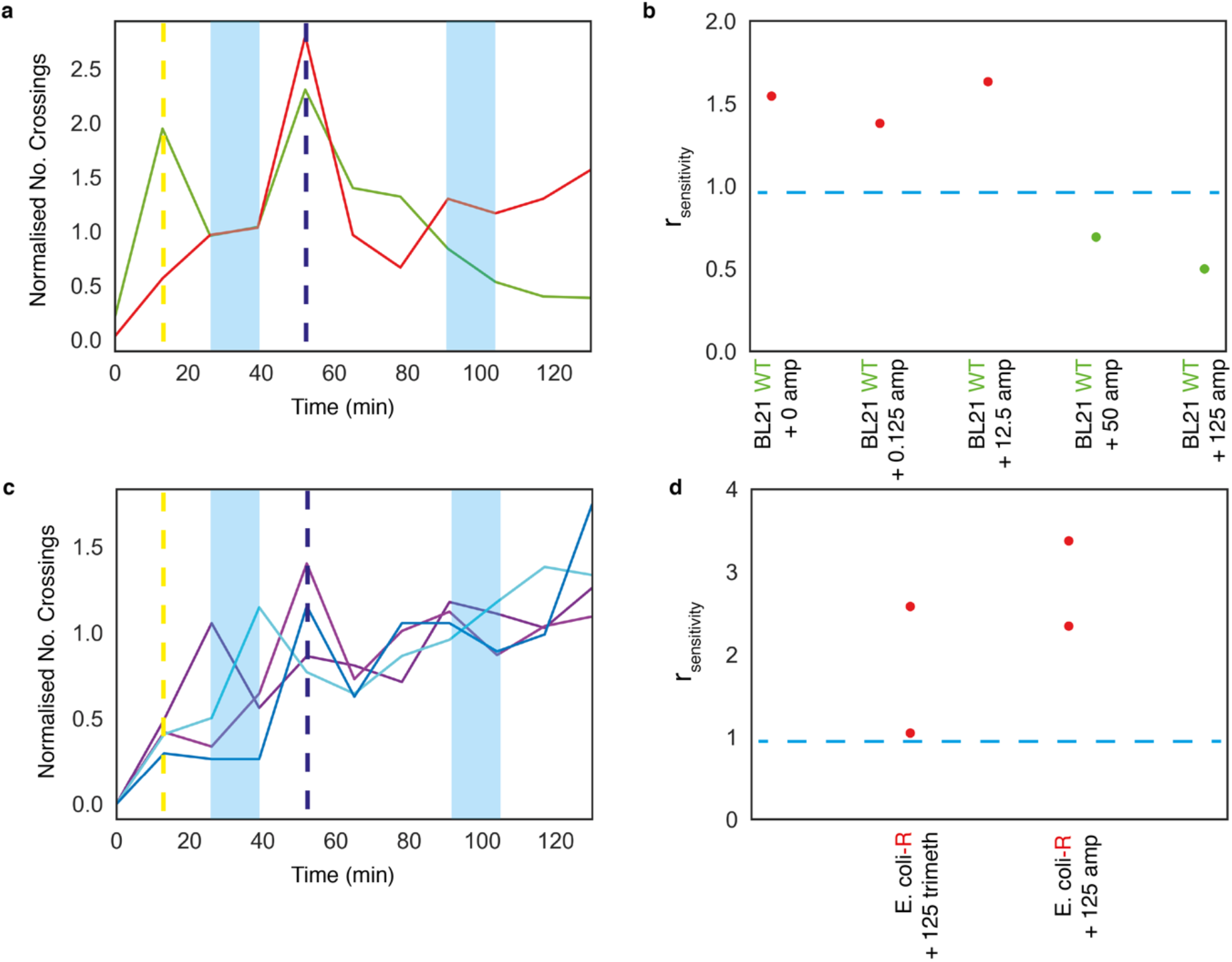
Systematic analysis of susceptibility in clinical and laboratory strains of *E. coli*. **a**, Susceptibility of BL21-WT (S, green) and BL21-ampR *E. coli* (R, red) to 125 μg/mL ampicillin. Addition of bacteria (yellow dotted line) and antibiotic solution (dark blue dotted line) to the system cause large fluctuations in the signal as the liquid is mixed, which dissipate within ~800 seconds. Number of bacterial crossings in a given time period, here 800 seconds, is plotted. The number of bacterial crossings shows a decrease 45 minutes after antibiotic addition. **b**, Determination of resistance profile, with sensitivity readout (r_sensitivity_). r_sensitivity_ was calculated using the ratio of crossings post-antibiotic and pre-antibiotic at set time points marked in blue in **a**. Strains were determined to be sensitive (S) if r_sensitivity_ < 1 (green); or resistant (R) if r_sensitivity_ ⩾ 1 (red), cut off (r_sensitivity_ = 1) shown as blue dashed line. Shown for five concentrations of ampicillin and BL21 *E. coli* **c**, Susceptibility of a clinical isolate of *E. coli*, determined to be resistant to both ampicillin (purple lines) and trimethoprim (blue lines). **d**, Determination of resistance profile. r_sensitivity_ for repeats of clinical isolate with 125 μg/mL trimethoprim and ampicillin. Antibiotic concentrations are given in μg/mL.

Using this method sensitive and resistant strains of *E. coli* can be differentiated. As described above, a reduction in signal after addition of antibiotic for sensitive strains is seen (Figure 3a, green); for resistant strains, there is an *increase* in signal (Figure 3a, red). Though the trend remains the same, the magnitude of the signal change can vary (Figure S6a) based on several factors which effect growth rates, including inoculant concentration, strain, and temperature, for example. The data was therefore normalized to the baseline taken before the addition of antibiotic when comparing between experiments (S_baseline_) (Figure S6b).

To obtain a systematic readout of antibiotic sensitivity across experiments, including multiple strains and antibiotics, a normalized measure of bacterial growth was determined as follows. Antibiotic sensitivity (r_sensitivity_) is defined as the ratio of S_baseline_ and 45 minutes post-antibiotic treatment (S_antibiotic_), shaded blue in Figure 3a. r_sensitivity_ provides a binary readout of sensitivity, r_sensitivity_ ≤ 1indicates cell death or inhibition of bacterial growth, and sensitivity to the antibiotic in solution; r_sensitivity_ > 1 indicates bacterial growth, and therefore resistance to the antibiotic used. This method allows for both bactericidal and bacteriostatic antibiotics to be used, as r_sensitivity_ < 1 indicates a decrease in cell number, or cell death (bactericidal); r_sensitivity_ = 1 would indicate inhibition of growth, but little cell death (bacteriostatic). For Figure 3a with ampicillin, r_sensitivity_ = 0.5 for the green strain (sensitive) and r_sensitivity_ = 1.1 for the red strain (resistant). For kanamycin, r_sensitivity_ = 0.92 for a sensitive strain and r_sensitivity_ = 2.0 for a resistant strain (green and red, respectively Figure S7).

Having shown that r_sensitivity_ can be used as a measure of bacterial sensitivity, this method was applied across a range of concentrations of ampicillin to determine the minimum inhibitory concentration (MIC) for the *E.coli* strain BL21 (Figure 3b). The MIC value is defined as the lowest concentration of an antibiotic that will inhibit the visible growth of a bacterial strain,^[21]^ and is used to inform clinical breakpoints and provide patient-dose information for prescribing treatment. At low ampicillin concentrations (0-12.5 μg/mL r_sensitivity_ > 1, and at increased ampicillin concentrations (50-125 μg/mL) r_sensitivity_ < 1. This indicates an MIC of 12.5-50 μg/mL ampicillin for *E.coli* BL21. This result is within the range determined by broth microdilution, the gold standard method (8-16 μg/mL). Despite difficulties in variability of measuring MICs,^[22, 23]^ these values are used by clinicians when making decisions about patient care (antibiotic selection and dosing), and hence are an important result for any new diagnostic tool to accurately measure.

Uropathogenic *E. coli* (UPEC) is the leading cause of urinary tract infections (UTIs),^[24]^ and is clinically burdensome across the globe. AMR has increased in UTIs and hence represents an excellent clinical target for a new diagnostic tool. The potential of this rapid optical AST method was demonstrated by testing on an *E. coli* clinical isolate. As shown in Figure 3c, treatment of the clinical isolate with 125 μg/mL ampicillin and trimethoprim resulted in no decrease in signal, and gave r_sensitivity_ > 1 within 45 minutes (Figure 3d). This was confirmed by broth microdilution (resistance >256 μg/mL ampicillin and trimethoprim). These detected resistances agreed with the resistance spectrum obtained from the hospital (Great Ormond Street Hospital, London) measured by the gold standard method in the clinical laboratory (Table S1). This study demonstrates the ability of this method to successfully carry out an AST for a strain of bacteria isolated from a patient within 45 minutes of the addition of antibiotic.

To conclude, in the face of AMR novel rapid methods to detect resistance in bacteria are needed to prevent its further spread and development. This study has shown that this rapid optical AST method can rapidly differentiate between resistant and sensitive phenotypes in lab and clinical strains of *E. coli* and determine MIC values to the same range as current gold standard methods. A read out of bacterial sensitivity was obtained within ~45 minutes of the addition of antibiotic. This method lends itself to miniaturization and automation, requiring a stable reflective surface which could be embedded within a 96-well plate for automated reading, with a laser and photodetector readout. This method can be exploited as a new rapid phenotypic method for AST, to provide these time-critical results to inform patient care and antibiotic stewardship.

### Experimental Section

#### Experimental method

A stiff AC160 TS cantilever (*k* = 26 N/m; Olympus, Japan) was loaded onto an AFM head (JPK Nanowizard 3 ULTRA Speed; JPK Instruments, Germany) and immersed in filtered Luria Broth (LB; Sigma-Aldrich, USA) in a 35 mm diameter glass bottom petri dish (WillCo Wells, Netherlands). The cantilever spring constant was calibrated using the thermal noise method in the JPK software to convert vertical deflection from volts to nm. The cantilever was allowed to equilibrate for 15 minutes, during which time vertical deflection of the laser was measured. The LB media was then inoculated with bacteria to a constant concentration (~10^5^ CFU) and recording was started again for another 40 minutes to obtain pre-antibiotic baseline. Antibiotic solution was then added to directly to the LB + bacteria solution to a desired final concentration, and deflection recording was then measured.

During experiments only the real-time scan function was used to monitor vertical deflection of the laser. Experiments were conducted at 28°C in an acoustic isolation hood. Prior to the start of the experiments, the AFM laser was left on for ~2 hours to ensure the laser had warmed up fully and to reduce laser power fluctuations which would affect the drift of the signal.

#### Reagents

Luria broth (LB) and antibiotics (ampicillin, kanamycin, trimethoprim) were all supplied by Sigma-Aldrich (USA).

#### Bacterial Strains

*E. coli* BL21(DE3)pLysS competent cells (Promega, UK) were selected for their suitability for transformation with a plasmid containing ampicillin resistance (pRSET/EmGFP plasmid; Invitrogen, UK).

A clinical isolate of *E. coli* was obtained from the microbiology repository of Great Ormond Street Hospital (London, UK).

#### Bacterial preparation

An LB media (Sigma-Aldrich) plate was streaked with BL21 *E. coli* (Promega) or clinical isolate *E*. coli (obtained from Great Ormond Street Hospital) from frozen stocks in a sterile hood. These were grown up overnight at 37°C. A single colony was used to inoculate 4 mL LB media, which was incubated at 37°C for 2 hours (225 r.p.m. shaking), to obtain mid-log phase growth. The OD600 of the culture was measured using a Nanodrop One-C (Thermo Scientific), and a final OD600 for bacterial inoculation for experimental measurement was adjusted to keep as constant as possible.

#### Bacterial transformation with ampicillin resistance

An aliquot of competent bacterial stock was thawed on ice for 20-30 minutes. 1-5 μL (10pg-100ng) pRSET-EmGFP plasmid (Invitrogen, CA, USA) was mixed with 25 μL thawed bacterial solution and incubated for 5-10 minutes on ice, followed by heat shock treatment at 42°C for 40 seconds and returned to ice for two further minutes. 500 μL warmed SOC media was added, and this was incubated at 37°C at 225 r.p.m. for one hour. 50 μL was plated onto an agar plate which contained 50 μg/mL nafcillin/ampicillin mixture. This plate was incubated overnight at 37°C and colonies used were made into frozen stocks for experimental use.

#### Data analysis

Vertical deflection data (nm) was recorded on JPK Nanowizard 3 software at 20 kHz sampling frequency. This raw data (Figure S8a) was then processed in 800 second “chunks” using analysis code written in Matlab. This code applies a Savitzky-Golay finite impulse response (FIR) smoothing filter of polynomial order 2 to the data, with a filtering frequency of 101 Hz (Figure S8b). A Savitzky-Golay smoothing filter was chosen as this function can filter noisy data effectively without removing high frequency data.

To identify the number of bacterial crossings, both local maxima and minima were identified, as bacteria moving through the laser was observed to cause both peaks and dips in the signal (Figure S8c, peaks labelled with blue triangles). A “Peak Finder” function was used to identify local minima/maxima in the signal, where a “peak” was defined as having a threshold drop of at least 0.5 nm on each side. This was to ensure that only the larger peaks were counted, which correspond to bacteria moving across the laser. Smaller “noise” seen in the signal was not attributed to actual bacterial crossings, but could be due to partial crossings, or a change of orientation of bacteria within the laser during a crossing. This threshold peak prominence value of 0.5 nm was applied empirically across all files when carrying out the analysis to remove any bias of identifying peaks in the signal.

Across the experiment, the number of peaks was calculated for a subsampled time frame to increase the resolution of the data from 800 seconds to 267 seconds, and plotted across the experimental conditions of LB media, addition of bacteria, addition of antibiotic (Figure S8d).

To calculate the antibiotic sensitivity (r_sensitivity_) the ratio of the signal pre-antibiotic addition, S_baseline_, and 45 minutes post-antibiotic addition, S_antibiotic_ (Figure S8d). r_sensitivity_ provides a binary readout of sensitivity, r_sensitivity_ ≤ 1 indicates cell death or inhibition of bacterial growth, and sensitivity to the antibiotic in solution; r_sensitivity_ > 1 indicates bacterial growth, and therefore resistance to the antibiotic used.

## Acknowledgements

This work was supported by i-sense EPSRC IRC in Early Warning Sensing Systems in Infectious Disease (EP/K031953/1), the European Metrology Programme for Innovation and Research (EMPIR) joint research project [HLT07] “AntiMicroResist” which has received funding from the EMPIR program co-financed by the Participating States and the European Union’s Horizon 2020 research and innovation program, i-sense EPSRC IRC in Agile Early Warning Sensing Systems for Infectious Diseases and Antimicrobial Resistance (EP/R00529X/1), EPSRC Royal Society Wolfson Research Merit Award, and by UKRI / MRC Rutherford Innovation Fellowship (MR/R024871/1). I.B. funded by EPSRC UCL Impact Award Grant affiliated to i-sense. The authors would like to thank E. Gray (UCL) for microbiology knowledge, K. Harris and R. Doyle (GOSH) for providing the clinical isolate, and T. Evans and B. Miller for assistance with data analysis (UCL).

I.B., A.L.B.P. and R.M.K. designed the study. I.B. performed experiments. I.B. and A.L.B.P. analyzed the data. I.B., A.L.B.P. drafted the paper. All authors discussed the results, and edited the manuscript.

