## Supplementary Information for "Cantilever Sensors for Rapid Optical Antimicrobial Sensitivity Testing"

**SI Methods: Replication of Nanomechanical method.**

Bacteria were cultured overnight at 37 ◦C, 250 r.p.m. (~$10^6$ CFU/mL). The bacterial suspension was then washed in PBS, where 1 mL of bacterial suspension was centrifuged at 5000 r.p.m. for 1 minute. The supernatant was removed and the bacterial pellet re-suspended in 1 mL of PBS. This was repeated three times. The final bacterial solution was concentrated by a factor of 8x or 20x.

25% glutaraldehyde solution was diluted down to 0.5% in DI water. A small droplet was placed over cantilevers B and D (B: $k = 0.12, f_{res} = 23$ kHz; or D: $k = 0.06, f_{res} = 4$ kHz) of a DNP-S1 or NP-O10 chip (Bruker, USA) and incubated for 10 minutes. This was washed carefully with DI water and allowed to dry. A droplet of the concentrated bacterial solution was incubated on the same side for 30 minutes. Loose bacteria were washed off by dipping the cantilever gently into a petri dish of PBS. Bacterial immobilization cover was checked using a bright field microscope.

Experiments were carried out on a JPK Nanowizard™ 3 ULTRA Speed AFM system (Bruker, USA) using DNP-S1 or NP-O10 cantilevers. The AFM was operated in contact mode for cantilever calibration. During experiments only the real-time scan function was used to monitor vertical deflection. Experiments were conducted at room temperature. Prior to the start of the experiments, the AFM laser was left on for ~12 hours to ensure the laser had warmed up fully and to reduce laser power fluctuations which would affect the drift of the signal.

Cantilevers were calibrated prior to bacterial immobilization. Sensitivity (nm/V) and spring constant ($k$) were measured by force curve analysis and thermal tune method, respectively. These were noted for conversion of raw mV data into nm.

The antibiotic, ampicillin, was used to kill the bacteria, and was added to a final concentration of 125 μg/mL (far above the minimum bactericidal concentration (MBC)).

**Supplementary Information Figures**

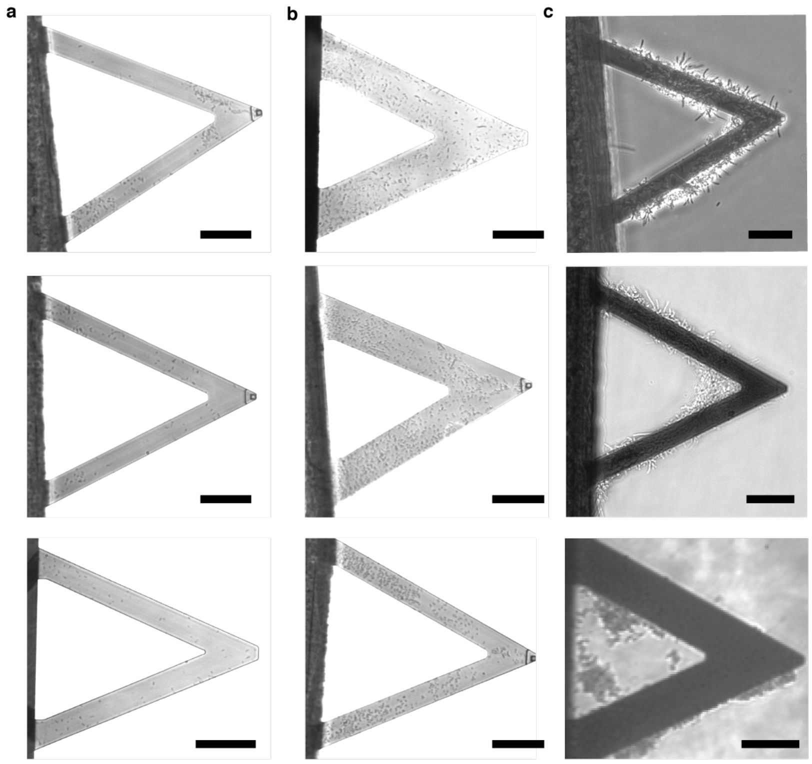


**Figure S1: Representative optical images of range in bacterial coverage.** a, Low bacterial coverage. b, “Optimal” (500-600 cells) bacterial coverage. c, high ‘clumpy’ bacterial coverage. Scale bars = 50 μm.

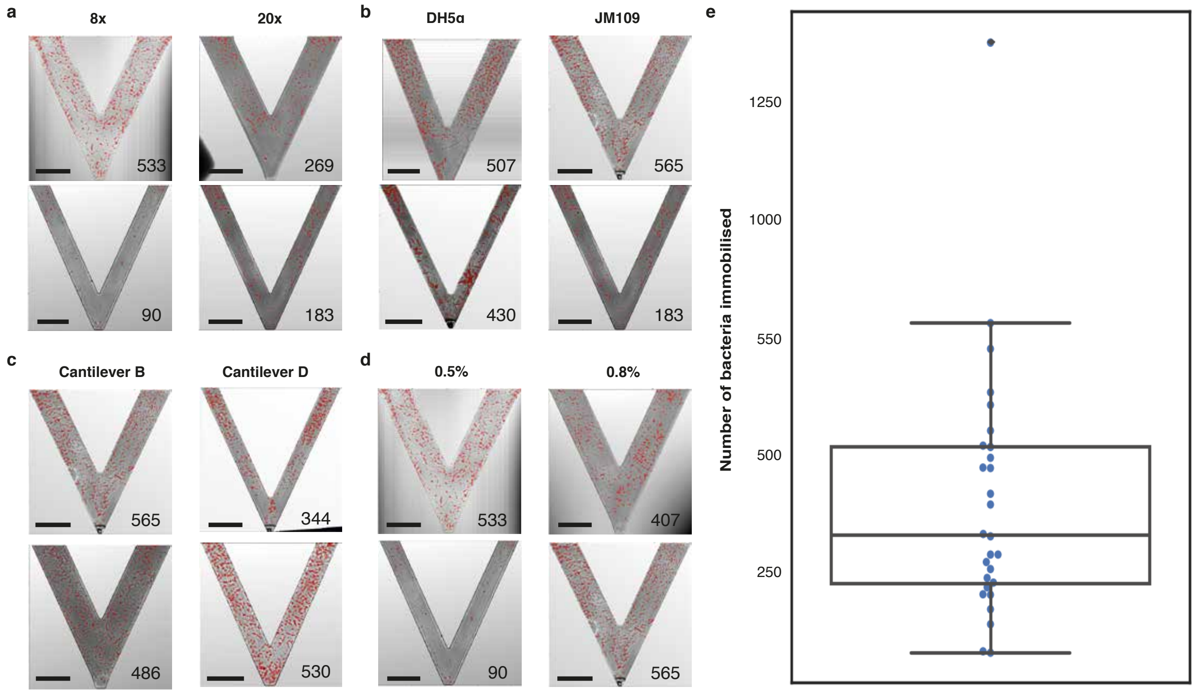


**Figure S2: Investigation of bacterial immobilization conditions.** Representative images of JM109 *E. coli*-immobilized cantilevers (DNP-S1 or tipless NP-O10), comparing different conditions and the number of bacteria immobilized. Other conditions are comparable across the cantilevers shown apart from the one specified. a, Cantilevers incubated with 8x (left) or 20x (right) concentrated bacterial solution for immobilization step. b, DH5α (left) and JM109 (right) used as the bacteria for immobilization. c, Comparison of immobilization levels on the greater surface area of cantilever B versus the narrower cantilever D. d, Cantilevers treated with 0.5% glutaraldehyde (left) or 0.8% glutaraldehyde (right) prior to immobilizing bacteria. Bacterial immobilization number estimate shown bottom right of each figure. Scale bars = 50 μm. e, Box plot showing spread of data points across 28 immobilization experiments. IQR across these is below 500 cells, which is below "optimal" immobilization level for nanomechanical experiments. Median = 342 cells; $Q_1$-$Q_3$ = 238-531 cells.

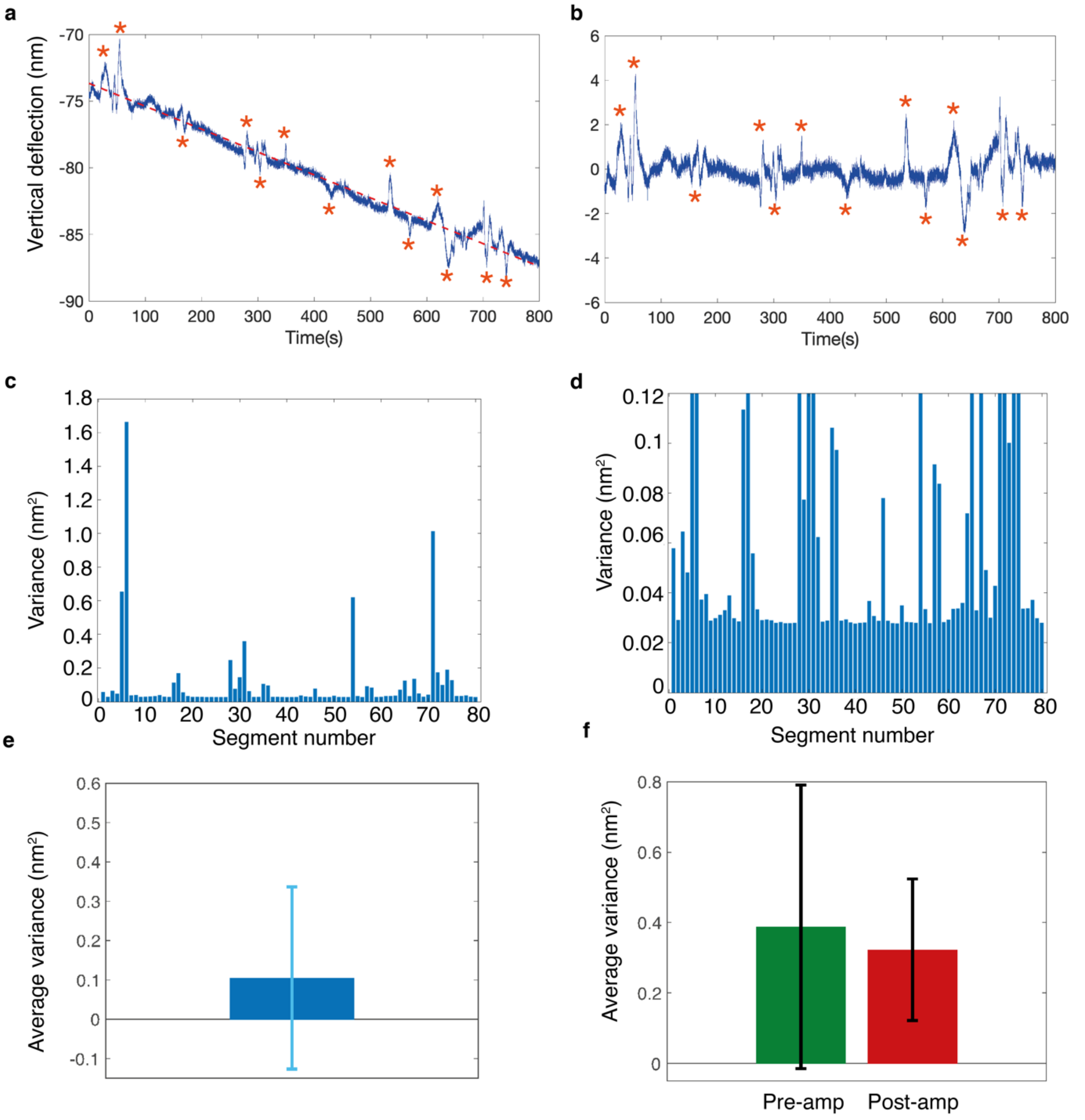


**Figure S3: Data analysis of mechanical signal. a-b**, Subtraction of linear regression from raw data, and large peaks not caused by mechanical motion of cantilever identified (*). **c-d**, averaging of variance over 10 second segments, removing large peaks from average variance calculation for one experiment (**e**). **f,** Average variance for n=5 experiments, pre- (green, pre-amp) and 15 minutes post-treatment (red, post-amp) with 125 µg/mL ampicillin for optimal immobilization count cantilever D experiments. Green box is pre-antibiotic treatment, red is post-antibiotic treatment. P=0.4569. Cantilever D: $k = 0.06$ N/m, $f_{res} = 4$ kHz.

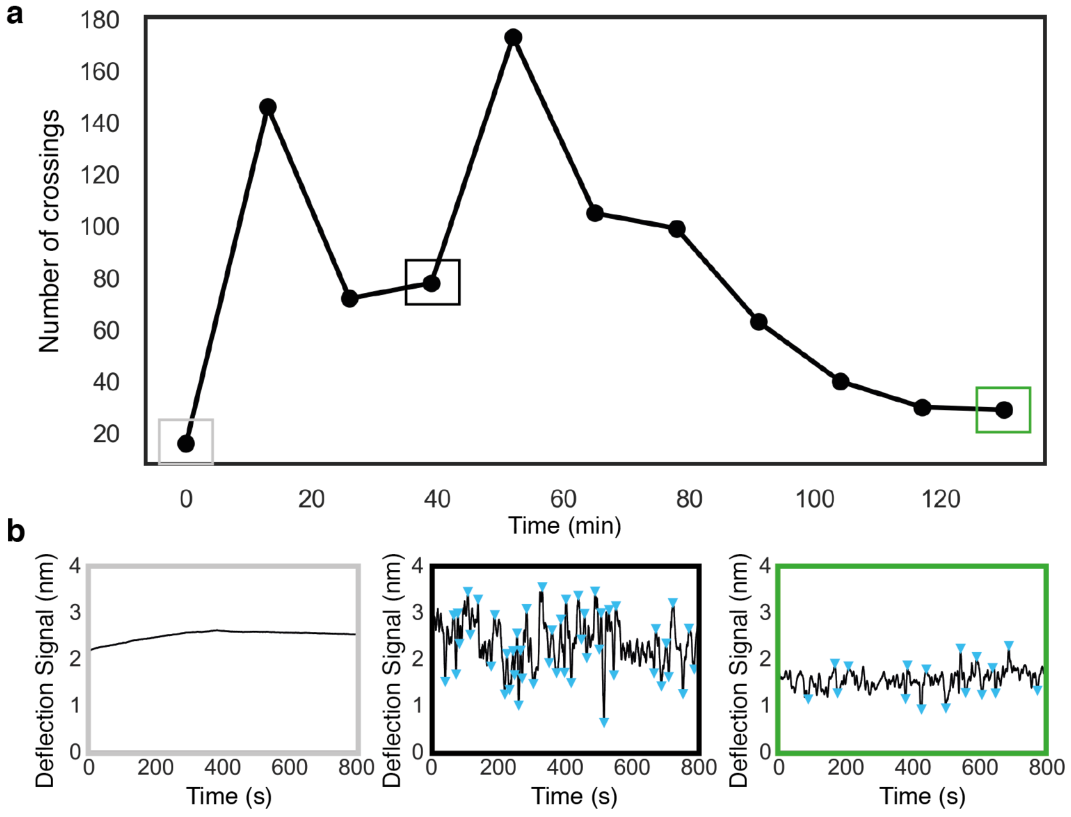


**Figure S4. Peak identifying and counting analysis. a**, The number of bacterial crossings in 800 seconds was calculated and plotted over course of experiment. **b**, Raw data traces for points at ‘media only’ (grey box), ‘inoculated media’ (black box), ‘inoculated media containing antibiotic’ (green box). Peaks identified (blue triangles) as ±0.5 nm from previous peak. Each point in **a** is total number of peaks identified in 800 seconds.

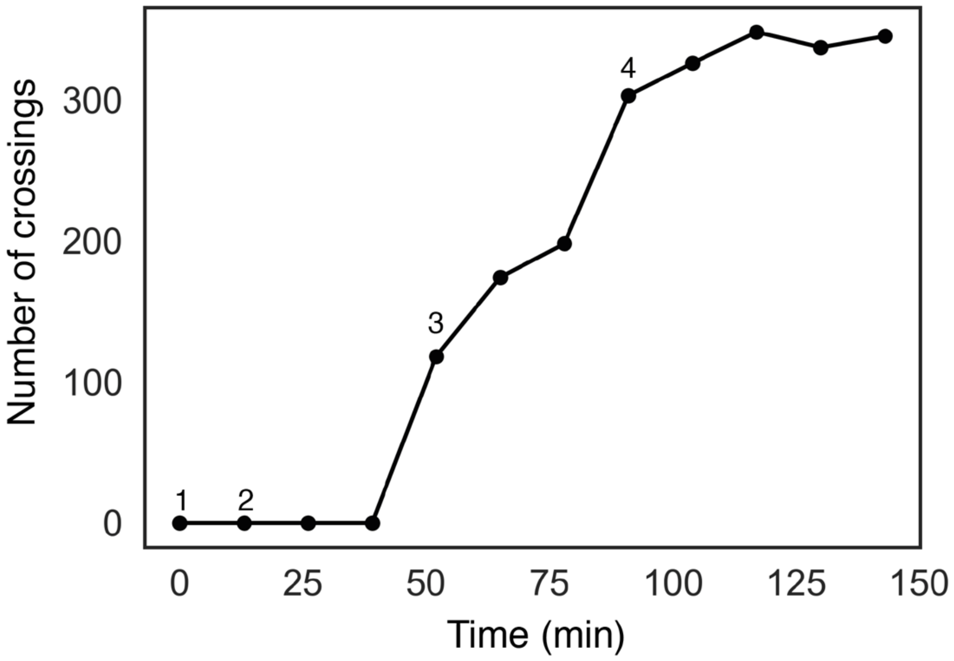


**Figure S5. Addition of media control.** Number of bacterial crossings in filtered media (1), addition of more filtered media (2), inoculation with BL21-WT *E. coli* cells, and addition of more filtered media (4).

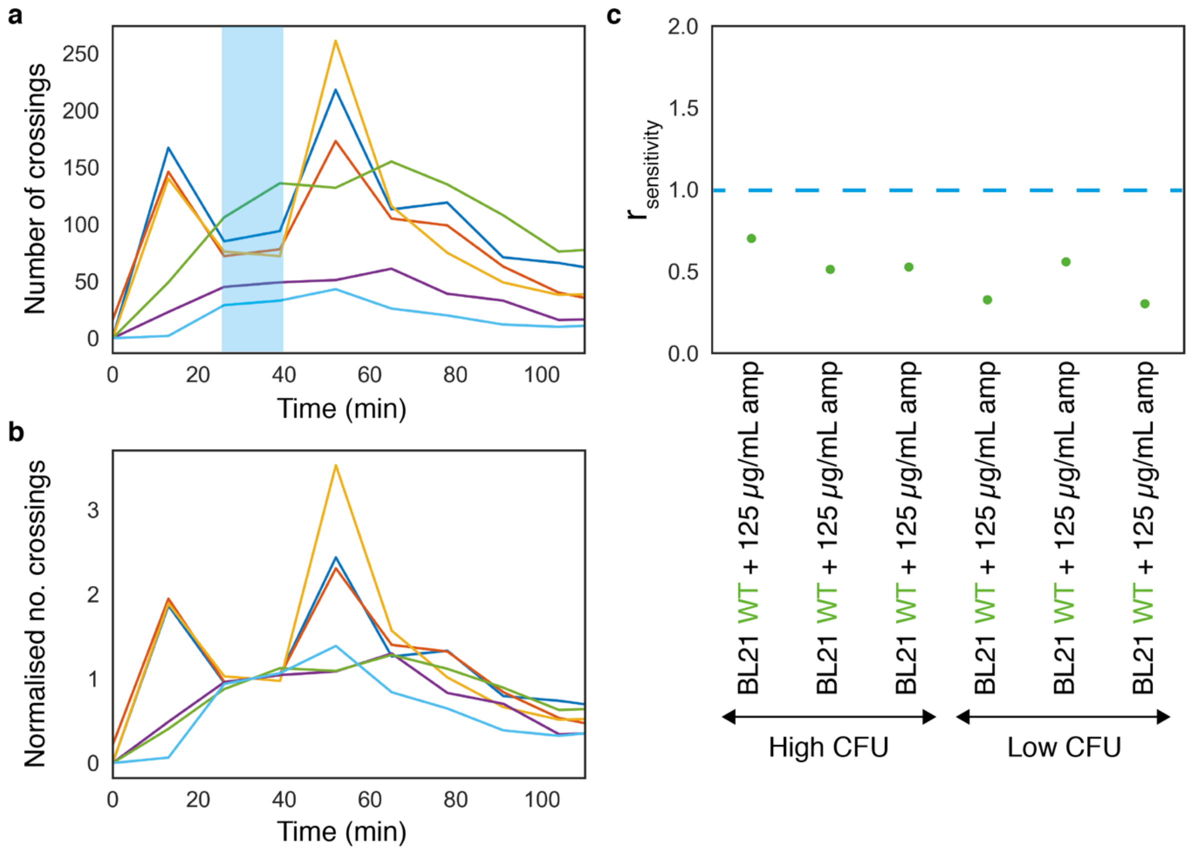


**Figure S6. Baseline normalization and magnitude variability between experiments. a**, Bacterial crossings over five experiments with BL21-WT *E. coli* (sensitive) and 125 μg/mL ampicillin. Yellow, dark blue and orange experiments were inoculated with a higher CFU of bacteria than purple and light blue experiments. **b**, Normalized data for same experiments as **a**. Normalization of data to baseline pre-antibiotic treatment (blue highlighted section) to remove variability in bacterial inoculation level so experimental data comparable across experiments. **c**, $r_{sensitivity}$ for five experiments shown in **a** and **b**. All show $r_{sensitivity} < 1$ (sensitive), demonstrating that initial inoculation level does not affect $r_{sensitivity}$ value.

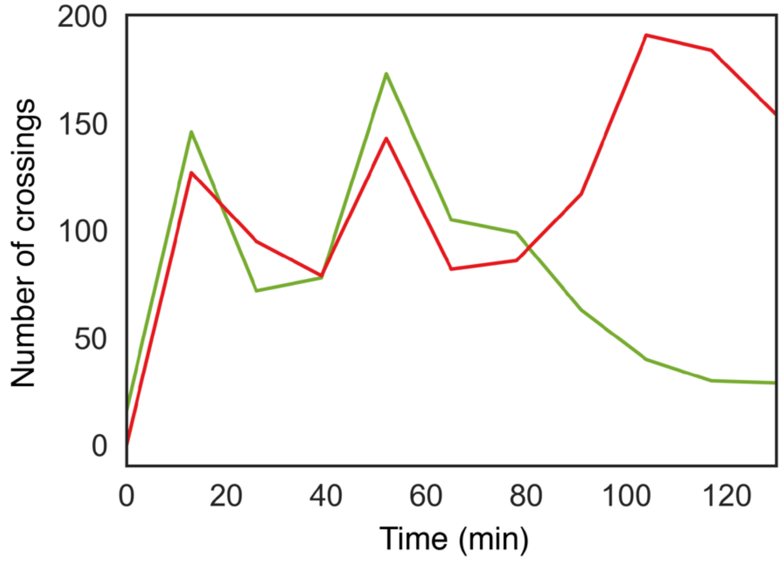


**Figure S7. Kanamycin resistant and sensitive strain.** Bacterial crossings over course of experiment for BL21-WT and BL21-kanaR + 125 µg/mL kanamycin. $r_{sensitivity} = 0.92$ for BL21-WT (sensitive strain, green) and $r_{sensitivity} = 2.0$ for BL21-kanaR (resistant strain, red).

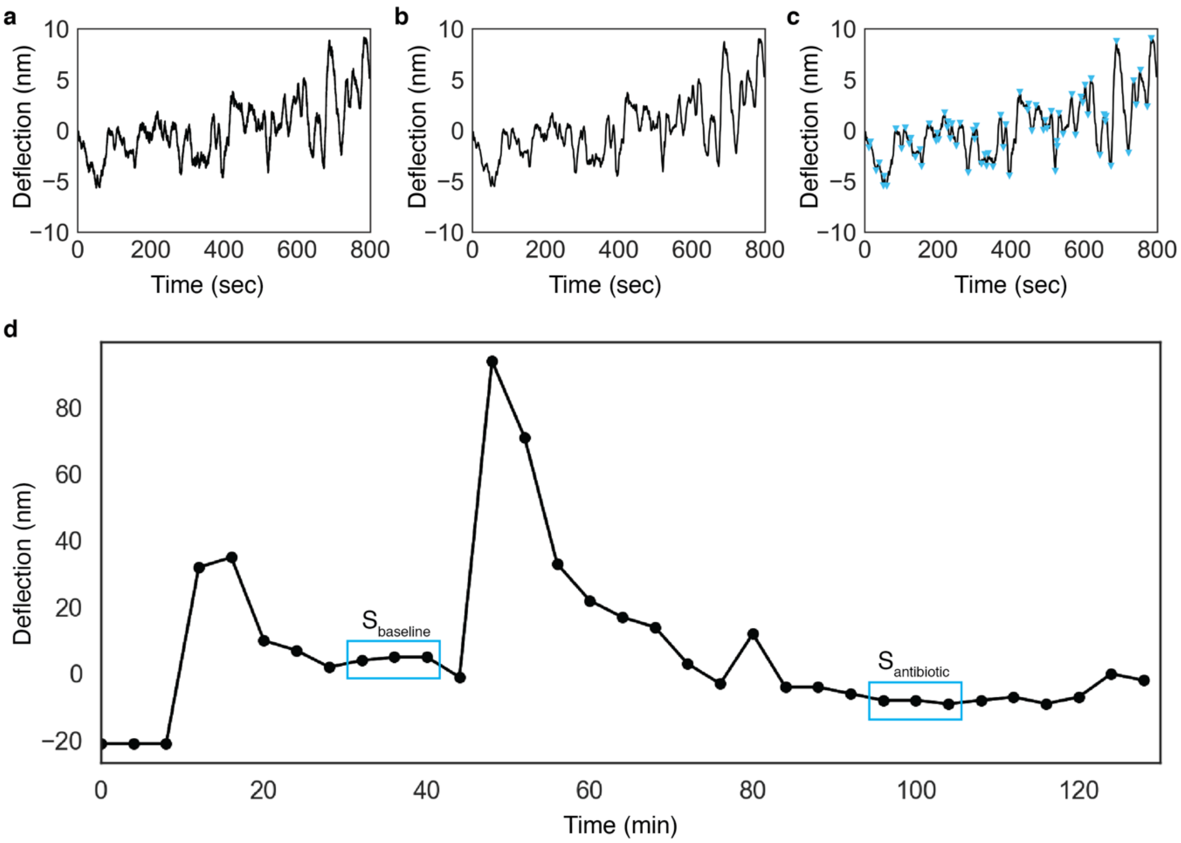


**Figure S8. Data analysis steps applied to raw data. a**, Raw data trace for vertical deflection taken over 800 seconds. **b**, Savitzky-Golay finite impulse response (FIR) smoothing filter of polynomial order 2 applied to the raw data, with a filtering frequency of 101 Hz. **c**, Peaks identified as having peak prominence value of 0.5 nm. This threshold was applied empirically across all files when carrying out the analysis to remove any bias of identifying peaks in the signal. **d**, Plot of bacterial crossings (number of peaks) across experiment. Each point is bacterial crossings collated for 267 seconds (800 seconds/3). $r_{sensitivity}$ is calculated by taking ratio of $S_{baseline}$ and 45 minutes post-antibiotic treatment, $S_{antibiotic}$. $r_{sensitivity}$ provides a binary readout of sensitivity, $r_{sensitivity} \leq 1$ indicates cell death or inhibition of bacterial growth, and sensitivity to the antibiotic in solution; $r_{sensitivity} > 1$ indicates bacterial growth, and therefore resistance to the antibiotic used. $r_{sensitivity}$ for this example was 0.66.

| Antibiotic | Resistant/Sensitive |
|---|---|
| Amikacin | R |
| Ampicillin | R |
| Augmentin | R |
| Cefotaxime | R |
| Ceftazidime | R |
| Chloramphenicol | R |
| Ciprofloxacin | R |
| Ertapenem | R |
| Fosfomycin | S |
| Gentamicin | R |
| Imipenem | S |
| Meropenem | R |
| Pipericillin/Tazobactam | R |
| Rifampicin | R |
| Temocillin | R |
| Tigecycline | S |
| Trimethroprim | R |
| Kanaymycin | R |

**Table S1. Resistance spectrum of patient isolate from Great Ormond Street Hospital.** Full resistance spectrum obtained using gold standard method for measuring break points for clinical isolate. Orange highlights two antibiotics chosen for this study, ampicillin and trimethoprim.
